# Sensor Fusion Algorithm to Improve Accuracy of Robotic Superposition Testing using 6-DOF Position Sensors

**DOI:** 10.1101/2024.12.16.627751

**Authors:** Callan M. Gillespie, Lesley R. Arant, Joshua D. Roth, Robb W. Colbrunn

## Abstract

To quantify the contributions of specific ligaments to overall joint biomechanics, the principle of superposition has been used for nearly 30 years. This principle relies on a robotic test system to move a biological joint to the same pose before and after transecting a ligament. The magnitude of the vector difference in joint forces before and after transecting the ligament is assumed to be the transected ligament’s tension. However, the robotic test system’s ability to accurately return the joint to the commanded pose is dependent on the compliance of the system’s various components, which is often neglected. An alternative approach to superposition testing is to use additional sensors attached directly to the joint to inform robot motion. Accordingly, there are two objectives in this manuscript: (1) **Describe** a testing methodology with 6-DOF position sensors to correct for system compliance, and (2) **Demonstrate** the effectiveness of this methodology to reduce uncertainty of in-situ forces determined using superposition. A Sensor Fusion algorithm fuses 6-DOF position sensors with robot pose measurements to compensate for system compliance. Using a surrogate knee joint with the Traditional testing method, errors in pose of approximately 100 microns resulted in a 23% underestimation of computed ligament tension. With the Sensor Fusion algorithm, errors in pose fell below the noise floor of the 6-DOF position sensors, and errors in computed ligament tension were reduced to 3%. Thus, this Sensor Fusion algorithm is a promising approach to minimize errors in superposition testing caused by compliance in a robotic test system.

## 1. INTRODUCTION

The use of robots to study the biomechanics of human cadaveric joints began in the early 1990s [1]. Just a few years after the early adoption of robots, the principle of superposition began to be applied to *in vitro* orthopedic biomechanical testing to quantify the contribution of ligaments to joint mechanics [2, 3]. Superposition testing is performed by applying a known overall load to a biological joint and recording the displacement of the joint at that load. After recording the displacement of the intact joint under the applied load, a structure within the specimen is then removed (e.g., cutting the Anterior Cruciate Ligament (ACL) in the knee).

Then, the joint is moved back to the recorded displacement, and a new load is recorded. This new load is the load carried by the remaining structures (e.g. everything but the ACL). The magnitude of the resulting vector difference between the original load and new load represents the load that was carried by the structure (e.g. ACL) that was removed [2].

In 1995, superposition testing was performed on a modified uniaxial testing machine with high stiffness [2, 3]. However, by the following year superposition testing began to be performed by 6-degree-of-freedom (DOF) serial arm robots, with inherently lower stiffness [4, 5]. The scope of these studies also expanded over the next several decades to include the use of the displacement measurements from these serial arm robots. The load-displacement relationship of various ligaments was examined in more detail and parameters such as *in-situ* ligament slack length and stiffness were also quantified [6–9]. Many of the ligament parameters elucidated by superposition testing are used by clinicians and medical device designers to

validate models, inform clinical understanding, and impact the design of new medical devices for joint replacement or repair [3, 6, 9–12].

One challenge with superposition testing is that the calculations assume that joint position is identical between different ligament conditions. In reality, the compliance of all components in the test system lead to systemic errors that, when unaccounted for, lead to underestimation of loads carried by the structures being tested [13]. All known robotic systems used for superposition testing at the time of writing this publication only control based on reported robot position as calculated by encoders within the robot’s joints. These encoders neither measure compliance of the individual links of the robot, nor measure compliance of the bones or other fixtures attached to the robot. Therefore these systems cannot differentiate displacements of the joint from displacements caused by compliance in other parts of the system [14]. These types of control systems are referred to as *Traditional* control systems herein. Traditional control systems rely on a Joint Stiffness Model assumption and so to remedy these shortcomings, a System Stiffness Model was proposed [13], where the compliance of each component of the system is stacked in sequence to account for the deflection of the robot, load cell, custom fixtures, and bones. In the System Stiffness Model, all components, other than the biological joint being measured, are lumped together and called the fixture stiffness. The System Stiffness Model predicted that superposition computed loads in the structure being tested have likely been underestimated in a Traditional control system because the joint has not necessarily been returned to the same position after the structure is cut. If system compliance could be compensated for, then displacement of the system could be dynamically corrected to return all structures within the joint to the same positions in all joint conditions. In turn, this correction enables the loads of any structure of interest within the joint to be accurately quantified.

There are two feasible methods to account for system compliance. The ***first*** method to account for system compliance and enforce consistent displacements between surgical states to accurately model all stiffness in the system to compensate for system deflection in real time.

However, characterizing stiffness in a 6-DOF test system such as a robot is difficult to accomplish because stiffness changes as a function of pose, payload, acceleration, and other parameters [15, 16]. A ***second*** method, proposed herein, relies on additional real-time kinematic measurements that are not influenced by system deflection. By attaching 6-DOF position sensors, such as optical motion capture markers, on either side of a biological joint, an independent measure of the position of each bone can be recorded in real-time and used by the system’s control algorithm to drive robot motion to match joint motion rather than the motion originally recorded by the robot. Closing the loop on a 6-DOF robotic test system with 6-DOF position sensors external to the robot’s encoders is a technique called visual servoing in the robotics field [17, 18]. Optical motion capture sensors were used as the 6-DOF position sensors in this study because they are readily available to biomechanists. In subsequent sections, the terms 6-DOF position sensor and motion capture sensor are used interchangeably. While this approach can correct for system compliance in real time, and thus has the ability to reduce compliance induced error, its performance is negatively impacted by the motion capture sensor’s low precision, low reliability, and latency.

Therefore, this paper introduces an alternative methodology wherein a Sensor Fusion algorithm utilizes the high precision, high bias [19], reliable, and responsive robot encoder data present in Traditional control methods and fuses this with the low precision, low bias, less reliable, and less responsive motion capture markers. By fusing these two sensor outputs together, a more accurate control system is made with both high precision and low bias. Such a system is able to dynamically transform both kinetics and kinematics to a control reference frame that is independent of system compliance. Accordingly, the objectives of this work are to: (1) Describe the methodology for compliance compensation utilizing 6-DOF motion capture markers and a Sensor Fusion algorithm, and (2) Demonstrate the effectiveness of this new superposition testing methodology to reduce uncertainty in in-situ force measurements in a surrogate knee model.

## 2. METHODS

### 2.1 Compliance Compensation

To understand how the Senor Fusion algorithm works, it is first necessary to understand how Traditional testing systems function. **Figure 1** provides a schematic of the Traditional control system kinematic chain used for such systems.

**Figure 1:**
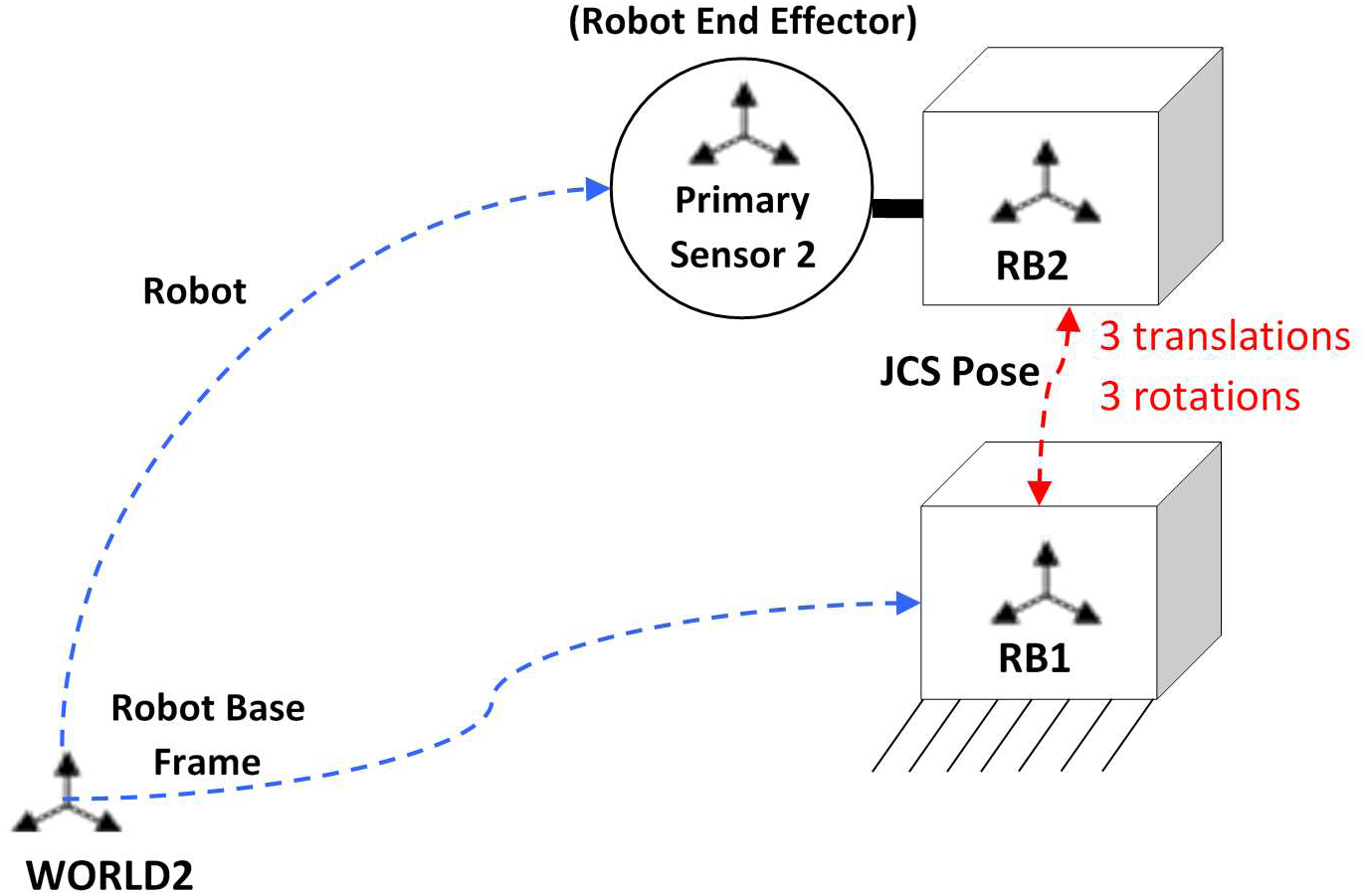
Schematic of the kinematic chain of a Traditional Testing System. This chain shows how the robot is the only sensor measuring joint motion. Displacement of the Rigid Body 2 (RB2 or bone 2) with respect to Rigid Body 1 (RB1 or bone 1) from its neutral position due to an applied load is considered its JCS (Joint Coordinate System) pose, which consists of 3 translations and 3 rotations. It is important to note that JCS pose here is the pose of the joint as reported by the robot, which means it does not compensate for fixture compliance.

In **Figure 1**, the concept of the Joint Coordinate System (JCS pose) is introduced, which describes the displacements of a joint, from its neutral pose, in 6-DOF in response to an applied load. In Traditional testing systems, JCS pose is computed solely based on reported robot position, which means it does not compensate for system compliance. To calculate JCS pose, 4×4 homogenous transformations matrices describing the relative relationships of various system components can be used. In transformation matrix form, JCS pose is the relative pose of RB2 with respect to RB1 (𝐓_𝐑𝐁𝟏_𝐑𝐁𝟐_) and can be described with the following kinematic chain:

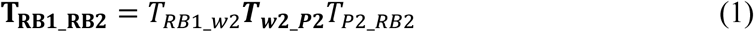

The two dynamically changing transformation matrices in the kinematic chain are the robot end effector pose relative to its base (𝐓_𝐰𝟐_𝐏𝟐_), and the relative pose of rigid body 2 relative to rigid body 1 (𝑻_𝑹𝑩𝟏_𝑹𝑩𝟐_). 𝑇_RB1_w2_ is the static relationship between rigid body 1 (RB1) and the robot’s base. Static relationships are defined by digitizing anatomic landmarks on the rigid body of interest as part of system setup. 𝑇_P2_RB2_ is the static relationship between rigid body 2 (RB2) and the Primary Sensor 2 (i.e., robot end effector pose). In superposition testing this is the relationship between the robot end effector and the bone attached to the robot.Importantly both 𝑇_P2_RB2_and 𝑇_RB1_w2_ are only static when there is no applied load on the system. As soon as there is an applied load then bones, custom fixtures, and the robot itself can deform, and this is not accounted for in the static relationships. After computing 𝐓_𝐑𝐁𝟏_𝐑𝐁𝟐_, the matrix is decomposed into 3 rotations and 3 translations according to International Society of Biomechanics Standard definitions for describing joint motion [20].

The Sensor Fusion approach (**Figure 2**) is designed to expect and measure system compliance in order to be able to compensate for it. This technique requires 6-DOF position sensors to be placed as close to the joint as possible to directly measure the joint’s pose. While the Sensor Fusion algorithm can work with any 6-DOF position sensor that can output data in real-time, optical motion capture sensors are focused on here as the 6-DOF position sensors because of their broad availability, familiarity, and utility in the biomechanics community. By placing motion capture sensors as close to the joint as possible, the measured joint pose will be less biased than the robot’s reported pose, which assumes that there is no compliance or displacement of the robot, fixtures, and bones while under load.

**Figure 2:**
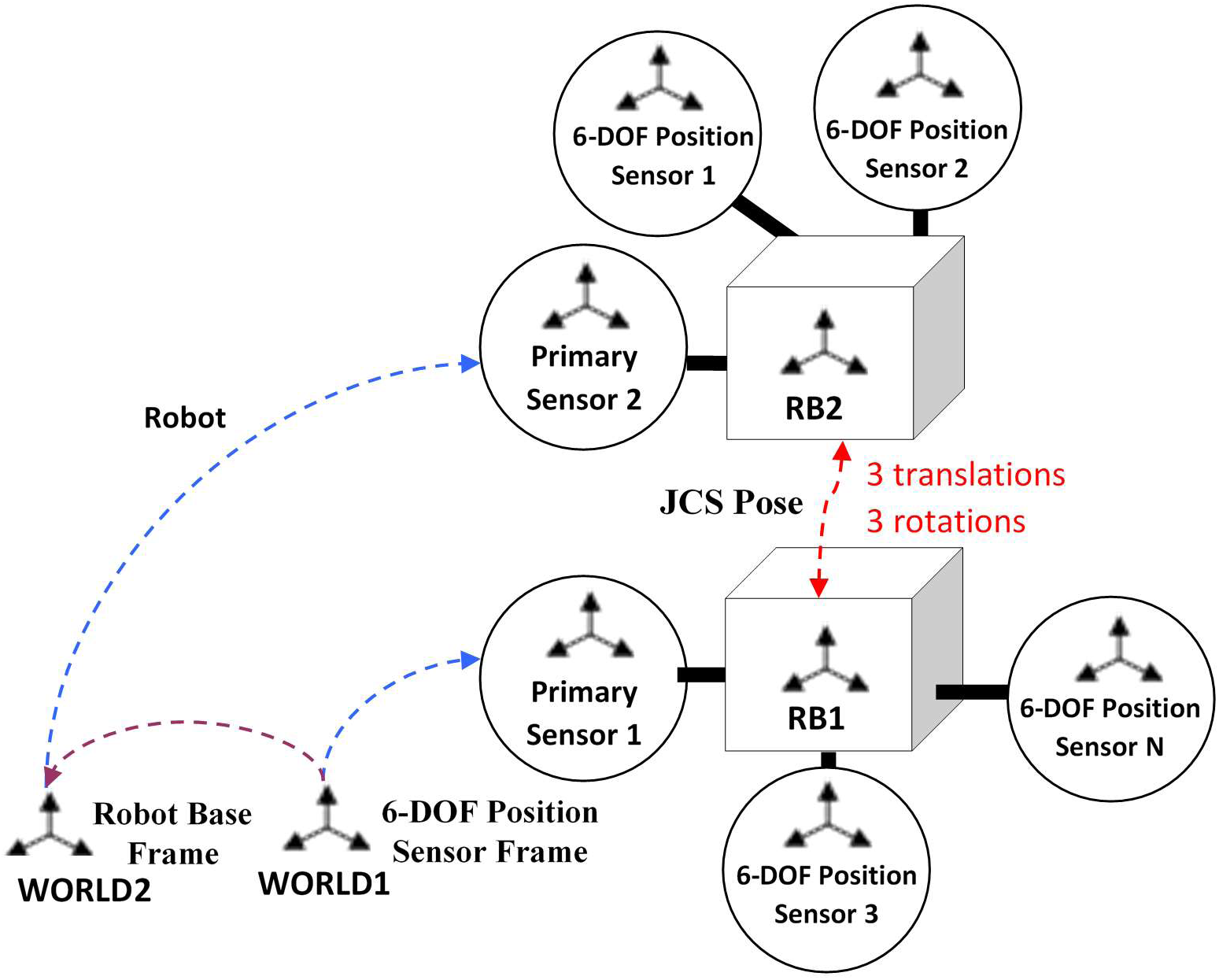
Schematic of the kinematic chain of a Sensor Fusion Testing System. This chain shows how the robot and additional 6-DOF position sensors both measure joint motion. Displacement of the Rigid Body 2 (RB2 or bone 2) with respect to Rigid Body 1 (RB1 or bone 1) from its neutral position due to an applied load is considered its JCS (Joint Coordinate System) pose, and it consists of 3 translations and 3 rotations. It is important to note that JCS pose here is the pose of the joint as reported by the robot *AND* 6-DOF position sensors, which means it does compensate for system compliance.

The Sensor Fusion algorithm permits the use of multiple motion capture sensors for each rigid body. During each control loop, the relative pose of the tool rigid body (RB2) must be calculated relative to the rigid body attached to the base (RB1), and the motion capture sensor data can be used to compute additional dynamically changing matrices in the kinematic chain.

These additional matrices (in bold) are dynamic and compensate for the compliance of the robot, fixtures, bones, and other structures in the testing system.

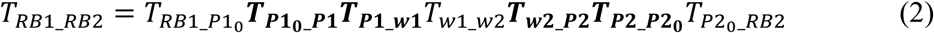

Unlike in (1), the base rigid body doesn’t have to be attached to a fixed structure in (2). The base could be an independently actuated or a passively moving structure and, in that case, would have its own sensor (Primary Sensor 1). 𝑻_𝑷𝟏_𝒘𝟏_is the dynamic relationship describing where the motion capture sensor frame (world 1) is relative to primary sensor 1 over time. 𝑇_RB1_P10_ is the static relationship between the base rigid body (RB1) and Primary Sensor 1 when first digitized.

𝑇_P20_RB2_ is the equivalent of 𝑇_P2_RB2_ from (1) with the only difference that it is explicitly indicated that it refers to where RB2 was when initially digitized. 𝑇_w1_w2_ is the relative spatial relationship of the robot base frame (world 2) with respect to the motion capture sensor frame (world 1).

𝑻_𝑷𝟏𝟎_𝑷𝟏_ describes the dynamic relationship between where the motion capture sensors calculate the Primary Sensor (**P1**) to be and where the Primary Sensor 1 was when first digitized (**P10**). This is considered the Delta 1 Matrix and is responsible for compensating for primary sensor 1, bone 1, or custom fixture compliance, collectively referred to as base compliance. If there are multiple motion capture sensors, then it is the quaternion weighted average of where they each calculate the Primary Sensor to be. 𝑻_𝑷𝟐_𝑷𝟐𝟎_ is the dynamic relationship between where the motion capture sensors calculate the Primary Sensor to be (**P2**) and where the Primary Sensor 2 was when first digitized (**P20**). This is considered the Delta 2 Matrix and is responsible for compensating for robot, bone 2, or custom fixture compliance, collectively referred to as tool compliance. Similar to the Delta 1 Matrix, it is a quaternion weighted average.

In equation (2), the bias of the calculated joint position is directly dependent on the noise, accuracy, latency, and placement of the motion capture sensors. This is why (2) is formulated in such a way that the Delta matrices are identity matrices unless there is compliance. The matrices represent deflection of the base and deflection of the tool rather than motion of the motion capture markers with respect to the camera system. Because system deflection should be lower magnitude and frequency content than marker motion, these matrices can be filtered more, further reducing noise. In other words, the robotic test system could be executing fast joint motion, but the Delta matrix changes will generally be significantly lower velocities.

In addition, the algorithm can accommodate multiple motion capture markers on a given rigid body to add redundancy to measurements. These 6-DOF motion capture data are combined with the Primary Sensor output via weighted quaternion averages to output a fused 6-DOF position in the form of a Delta Matrix. In the case where the motion capture markers are weighted at 100% and the Primary Sensor is weighted at 0%, the Primary Sensor output is not present in the Delta Matrix. However, the Primary Sensor will still be used in the sensor fusion algorithm shown in (2). For example, 𝑻_𝒘𝟐_𝑷𝟐_(robot position) is always present in (2), but corrected by the delta matrix 𝑻_𝑷𝟐_𝑷𝟐𝟎_ . In addition to manually weighting sensors, both the Primary and motion capture sensors can be weighted in an automated way based on reliability, proximity, Kalman filtering, or other models to further increase accuracy of the Delta matrices. Gravity compensation, load transformation, and even the position of the load cell itself can also be corrected by Delta matrices in kinematic chains similar to equation (2).

Most importantly, safety parameters are included in the Sensor Fusion algorithm. Passive markers might be mislabeled by a tracking system, motion capture markers may be occluded, or a marker might suddenly come back into view causing an instantaneous kinematic correction. To catch these scenarios, there are various kinematic safety parameters continuously monitoring the kinematic relationships of the motion capture markers both to each other and the Primary Sensor. If a marker is occluded, then its last known relative position is used until it comes back into view. When an occluded marker comes back into view, a secondary filter is applied to smoothly bring its output back into the control loop rather than instantaneously inject the output. Further, 6-DOF motion capture data can be interpolated via linear interpolation, but also via spherical linear interpolation, or other geometrically derived methodologies [21]. For these reasons, the Sensor Fusion method includes a sensor integrity handler that provides safety checks to reduce the risk of erroneous robot motion. The Sensor Fusion algorithm that performs compliance compensation with a sensor integrity handler to make the system safe is collectively called eXactoPOSE^TM^ More details on equation (2) and the integrity handler, are included in US Patent #11,745,341 B2 [14, 22].

### 2.2 Validation of Compliance Compensation

To directly measure how much ligament load is underestimated using the Traditional control method, and to validate the accuracy of the Sensor Fusion method, a ligament phantom spanning a surrogate knee joint was utilized (**Figure 3**) [23]. In subsequent sections of the paper it will be assumed that the structure being tested via superposition is this phantom ligament, and loads will be tension rather than compression. The surrogate knee contained 4 springs that had an equivalent spring stiffness of 354 N/mm in compression. The ligament phantom was positioned to simulate the function of the lateral collateral ligament (LCL). A single axis load cell (Futek LCM300) mounted on a spherical rod end pin directly measured ligament tension in the ligament phantom during superposition testing. Similarly, a motion capture system (OptiTrack, Corvalis Oregon) tracked 6-DOF reflective motion capture markers using Optitrak Prime^X^ 13 cameras placed close to the surrogate knee joint. These measurements were collected at the same time as the 6-DOF kinetics & kinematics measured by a 6-axis force/torque sensor (Omega160, ATI, Apex, NC) and a 6-DOF robot (KR 300 Ultra 2700-2, KUKA, Augsburg, Germany), respectively. The control load cell was mounted to the base. Joint motion was calculated using the Grood and Suntay coordinate system while joint loads were projected into the same non-orthogonal coordinate system [14, 24, 25].

**Figure 3:**
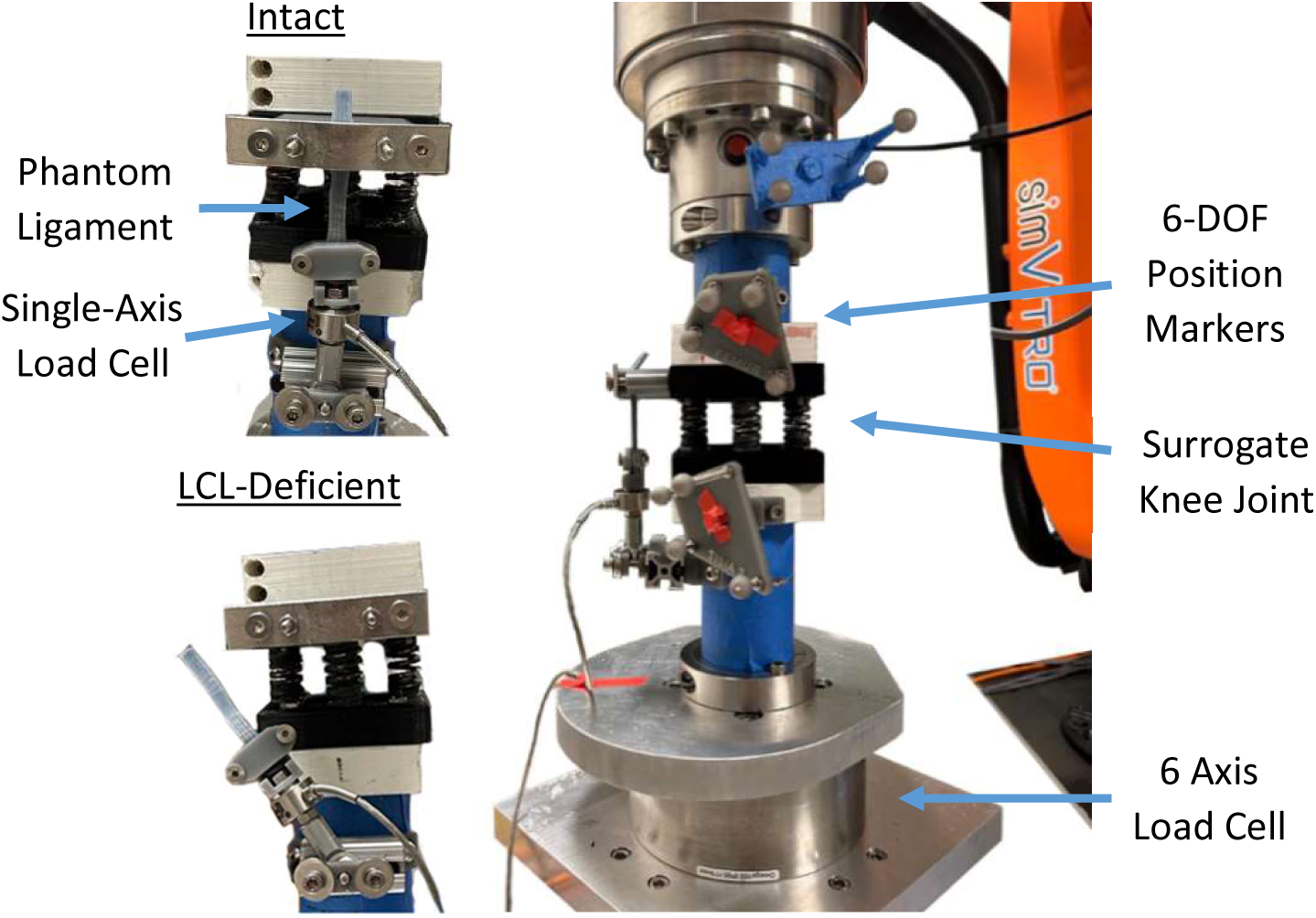
Composite image shows the surrogate knee attached to the robotic system. The surrogate knee joint was tested with an LCL phantom (intact) and without the phantom (LCL- deficient). LCL phantom tension was measured by both the single and 6 axis load cells.

To generate the initial kinematics for the superposition testing, a 6-DOF force/torque control test was performed independently by personnel from Biomechanical Advances in Medicine Lab at the University of Wisconsin-Madison using a simVITRO^®^ robotic test system (Cleveland Clinic, Cleveland, OH). The surrogate knee joint with the LCL phantom (**Figure 3**) was both loaded and unloaded with a 15Nm ramp of varus torque over 20 seconds, while prescribing a constant compression force of 50N. The peak 15 Nm varus torque was maintained for 5 seconds. All other DOFs were initially in force control and prescribed 0 N or 0 Nm, except flexion angle, which was in kinematic control and prescribed to be 0 degrees (**Figure 4**, Varus Force Control Test Block). This loading protocol was used to maximize tension in the LCL phantom during testing. The 6-DOF forces & torques were controlled in the JCS reference frame, and 2 sets of JCS pose kinematics were recorded simultaneously using both the Traditional technique and the Sensor Fusion technique. The recorded JCS pose kinematics were then low-pass filtered to reduce noise injected into the control signal by the motion capture markers.

**Figure 4:**
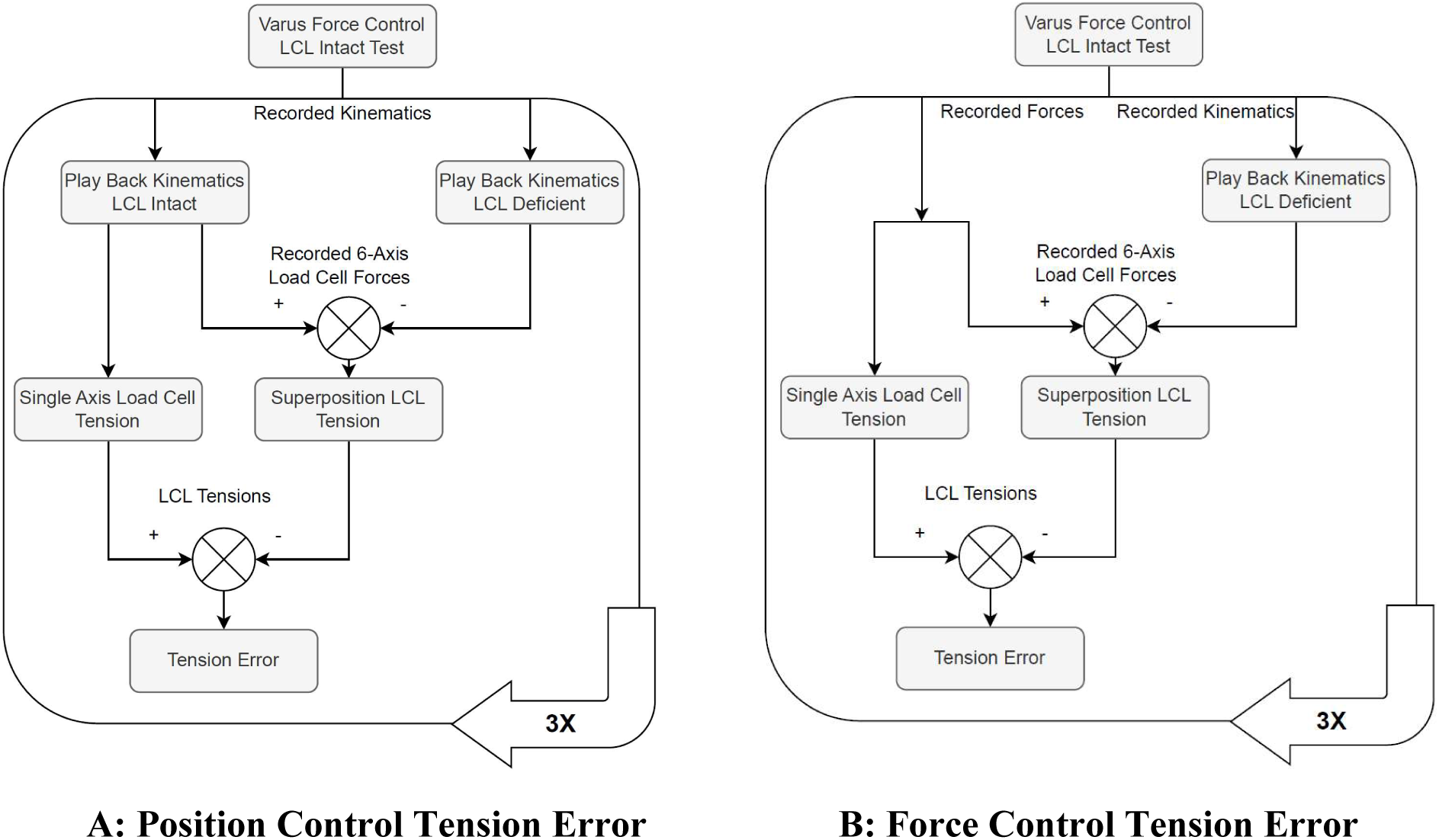
A flowchart of the testing completed to calculate the superposition LCL phantom Tension and Error. An initial varus force control test was performed, and robot and motion capture kinematics were recorded. In A, both these sets of kinematics were replayed 3 times with the LCL intact using Traditional control and then using Sensor Fusion Control. In B, the kinetics from the initial varus force control test were used. In both A and B, the LCL was removed, and both these sets of kinematics were replayed 3 times again.

As depicted in the flowchart in **Figure 4**, the recorded JCS pose kinematics were then played back using Traditional or Sensor Fusion control while also varying whether the surrogate knee had the LCL phantom intact (i.e., intact surrogate) or had the LCL phantom removed (i.e., deficient surrogate). The intact surrogate was tested 3 times in 6-DOF position control using Traditional control while prescribing JCS pose kinematics recorded using Traditional Control.

Similarly, the Sensor Fusion JCS recorded kinematics were played back on the intact surrogate 3 times while using the eXactoPOSE^TM^ algorithm to control kinematics. Then, the LCL phantom was removed, and the same tests were performed on the deficient surrogate. Throughout intact surrogate testing, the single axis load cell directly measured ligament phantom tension so it could be used for comparison with the superposition-computed tension in the ligament phantom. All recorded 6-DOF kinematics and kinetics were low-pass filtered at 0.2 Hz.

### 2.3 Analysis

*2.3.1 Tension Error at Peak Load*

LCL superposition-computed tension was computed using two approaches. In the first approach, which is most commonly used, the LCL superposition-computed tension (𝐿𝐶𝐿_SP,pos_) was calculated as the magnitude of the vector difference between intact 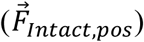 and deficient 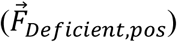 forces measured by the 6-axis load cell during the position control replays (**Figures 4** and **5A)** [2].

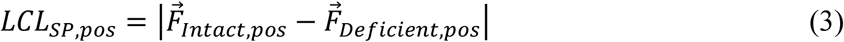

**Figure 5:**
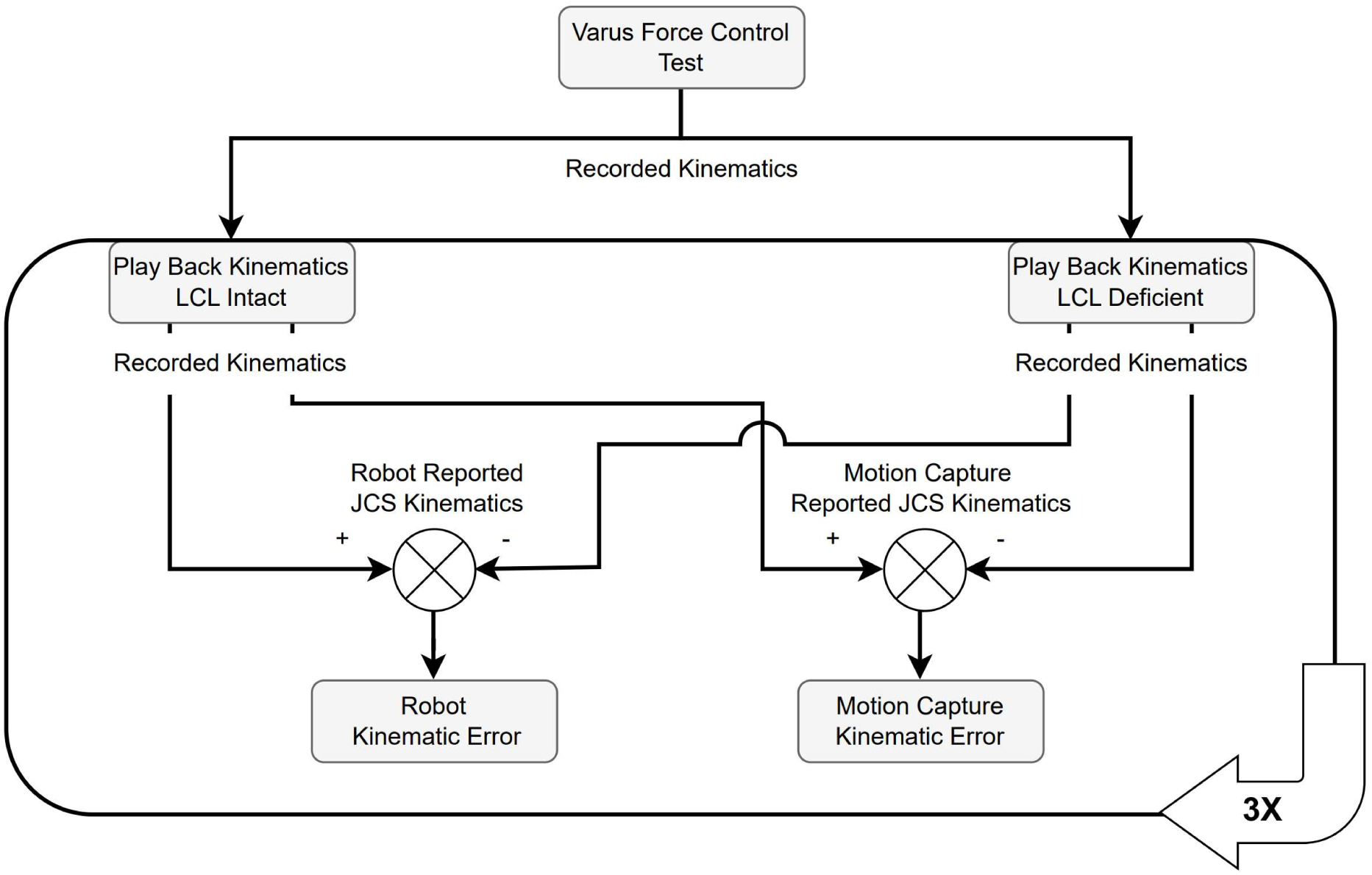
Calculation of Kinematic error. Both Traditional and Sensor Fusion kinematics were recorded during the Varus force control test and played back separately.

Tension error was calculated as the scalar difference between the tension directly measured by the Single Axis (SA) load cell in line with the LCL phantom (𝐿𝐶𝐿_SA,pos_) and the superposition-computed tension (𝐿𝐶𝐿_SP,pos_).

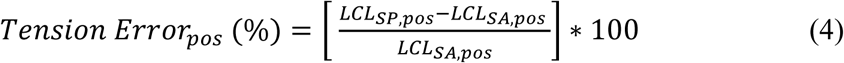

Tension error at peak load was computed as the average error at peak load, which was defined as the middle 3 seconds of the plateau at 15 Nm of varus torque. Tension error from each of the 3 trials was then pooled to compute a mean and standard deviation. Peak tension errors using this approach are termed the *Position control tension errors*.

A second approach was evaluated to determine whether it is possible to skip the intact position control test (**Figure 4B**) and assume that the positions measured during the original intact force control test are equivalent to the positions measured in the deficient position control test. In the second approach, the LCL superposition-computed tension (𝐿𝐶𝐿_SP,force_) was calculated as the magnitude of the vector difference between intact forces 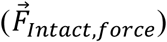 measured by the 6-axis load cell during the initial, varus force control trajectory and deficient forces 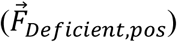 measured by the 6-axis load cell during the position control replay after sectioning the LCL (**Figures 4** and **5B)**. The tension errors were computed using the same equations as the first approach with the exception of substituting 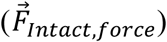 for 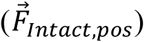, 𝐿𝐶𝐿_SP,force_ for 𝐿𝐶𝐿_SP,pos_and 𝐿𝐶𝐿_SA,force_ for 𝐿𝐶𝐿_SA,pos_. Peak errors using this second approach are termed the *Force control tension errors*.

*2.3.2 Kinematic Error at Peak Load*

Kinematic error 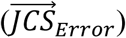 was calculated as the difference between the intact JCS pose 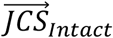 and deficient JCS pose 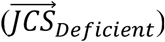.

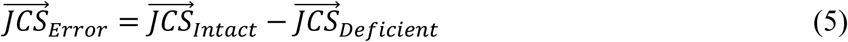

There were two characterizations of kinematic error for all superposition tests (**Figure 5**)

because joint kinematics were measured by both the robot and the motion capture sensors. Robot reported joint kinematics assume that the robot, fixtures, and bones do not have compliance. The motion capture kinematics do not make this assumption and therefore accounts for this compliance.

Equation (5) was useful to isolate kinematic error in a particular degree of freedom.

However, a scalar value representing resultant translation error was also analyzed to gain insight into control of translations between the two control methodologies. This error was calculated as the magnitude of the vector difference in the translation component of JCS pose.

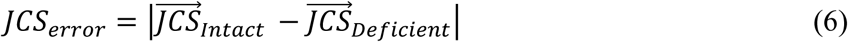

𝐽𝐶𝑆_error_ in (6) is the resultant distance between rigid body 1 and rigid body 2 as depicted by the translation components of JCS pose in **Figures 2** & **3**.

While it is helpful to understand translation error in the control reference frame (JCS), this coordinate system is not entirely indicative of the LCL phantom ligament length. Any error in JCS pose rotation will inject error into the stretch of the phantom ligament. As such, the LCL phantom ligament insertion points themselves were digitized, and their relative relationship was calculated throughout all kinematic trajectories. With or without the LCL phantom being intact, the insertion points can be used to calculate the LCL phantom length between insertions, which is an alternative metric for kinematic tracking relevant to ligament superposition testing.

Equation (6) was used a second time to calculate LCL phantom length error. This error metric was used because it is independent of the kinematic coupling of rotation and translation in the control reference frame.

As mentioned in **2.2 Validation of Compliance Compensation** , there were 2 superposition tests performed, the first was controlled by Traditional methodology and the other used Sensor Fusion. Therefore, Traditional and Sensor Fusion kinematic error at peak load was calculated as the average of kinematic error during the middle 3 seconds of the plateau at 15 Nm of varus torque. Kinematic error from each of the 3 trials was then averaged to compute a mean and standard deviation.

*2.3.3 Tension and Kinematic Error in Dynamic Loading*

While ligament contribution to joint stability (i.e. in situ loads) have historically been derived from superposition-computed tension at peak load, more recent work has examined ligament’s load-displacement relationship throughout the ligament’s engagement. Therefore, the tension and kinematic error when using Traditional and Sensor Fusion methodologies throughout dynamic loading is also of importance.

Initially RMS kinematic error between intact and deficient LCL phantom ligament length was calculated during ramping the varus torque up and down. Kinematic errors were so small that they cannot be easily seen when plotting any of the trial kinematic data. RMS tension error was not calculated because it can be visually seen in trial tension data.

The relationship between tension and error was also examined. It was hypothesized that as load applied to the test system increased then system deflection would increase, and as a result so would tension and kinematic error. To this end, both tension error and ligament length error were plotted against LCL tension as measured by the single axis load cell seen in **Figure 3**.

*2.3.4 Testing Control*

The kinematic control trajectories’ *control* was quantified for resultant translations and varus rotation by calculating the root mean squared error (RMSE) between the desired and actual kinematic setpoints. Resultant translation and varus rotation were focused on because these kinematic variables were most responsible for LCL engagement. For each control method the difference between desired and actual kinematic setpoints were pooled across the intact and deficient cases of the three superposition trials prior to running the RMSE. This metric quantifies the control system performance because a poorly tuned control system can introduce undesirable variability in the data and obscure the metrics of interest.

## 3. RESULTS

### 3.1 Tension Error at Peak Load

The Sensor Fusion control method underestimated ligament tension by substantially less than the Traditional control method using either of the two approaches to calculate tension error (**Figure 6**, **Table 1**). Using the Traditional control method, both the position control tension error and the force control tension error underestimated tension by 23%. Using the Sensor Fusion

**Figure 6:**
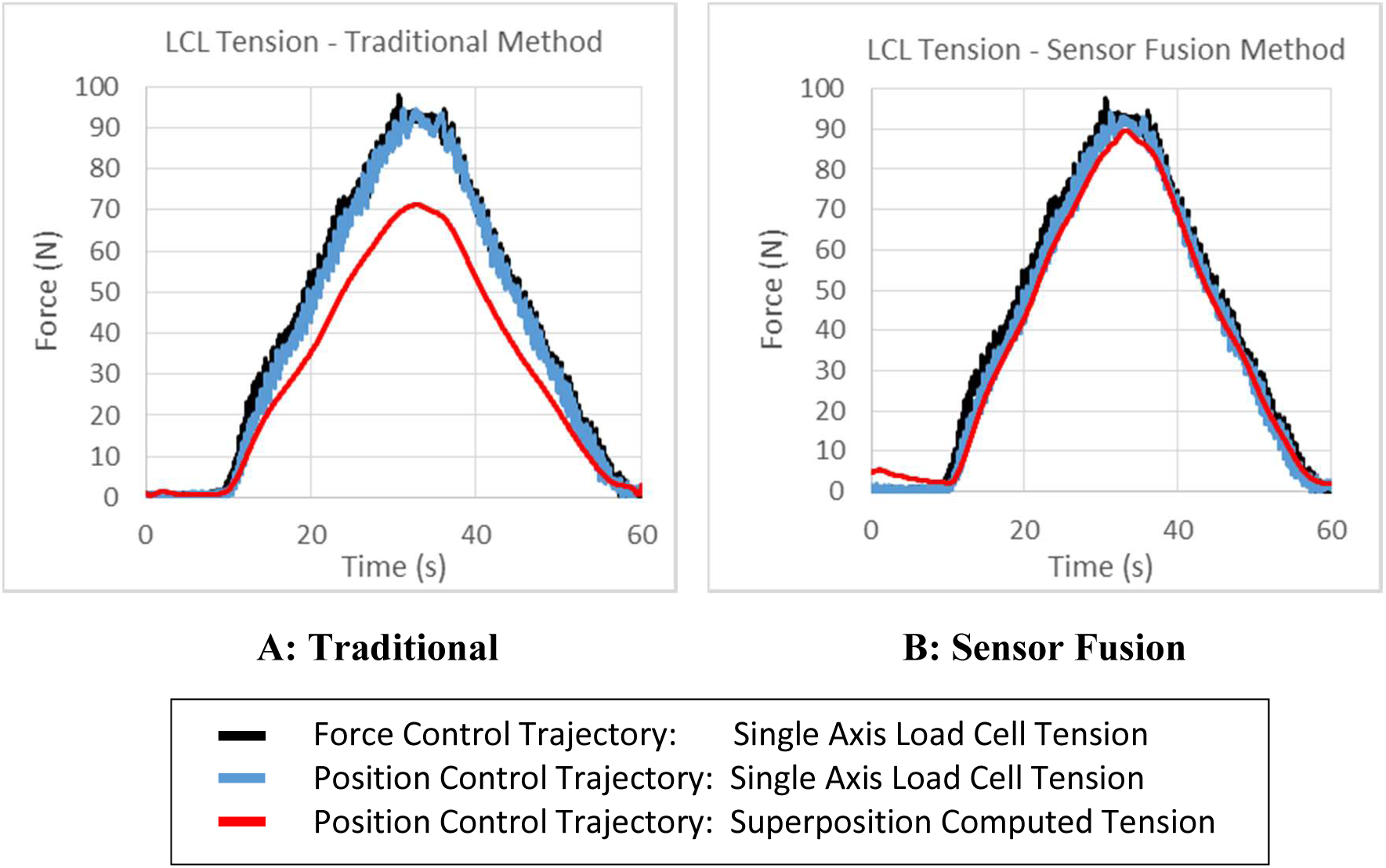
Line plots show the superposition-computed LCL phantom tension using Traditional and Sensor Fusion methods for control. Blue lines are the direct measure of tension in the LCL phantom by the single axis load cell. Red lines are what the superposition method computed the tension to be. A low pass filter was applied to the superposition-computed ligament tension data to smooth out the motion capture noise. This is why both Traditional and Sensor Fusion superposition-computed tension noise (red lines) look similar in Figure 6. The black line is the direct measure of tension during the initial varus force control test using Sensor Fusion for control as depicted in Figure 4.

**Table 1:** Tension errors at Peak Load using the Sensor Fusion method compared to the Traditional Method. These errors were computed using two approaches as the *position control tension error* and *force control tension error*. All data are reported as Mean ± SD of the percent error, which is normalized to the peak force.

| Control Methodology | Position control tension error (%) | Force control tension error (%) |
| --- | --- | --- |
| Traditional | -23 ± 0 | -23 ± 0 |
| Sensor Fusion | -3 ± 0 | -5 ± 0 |

**Table 2:** Mean ± SD of the Resultant Surrogate Translation Error between LCL Phantom Intact and Deficient Conditions at Peak Load.

| Control Methodology | Robot Reported Error ($\mu\text{m}$) | Motion Capture Reported Error ($\mu\text{m}$) |
| --- | --- | --- |
| Traditional | $<1 \pm 0$ | $85 \pm 1$ |
| Sensor Fusion | $100 \pm 1$ | $3 \pm 0$ |

**Table 3:** Mean ± SD of the Varus Surrogate Rotation Error between LCL Phantom Intact and Deficient Conditions at Peak Load.

| Control Methodology | Robot Reported Error (Deg) | Motion Capture Reported Error (Deg) |
| --- | --- | --- |
| Traditional | $<0.001 \pm 0$ | $-0.013 \pm .001$ |
| Sensor Fusion | $0.015 \pm .001$ | $<0.001 \pm 0$ |

control method, position control tension error was underestimated by 3%, while force control tension error was 5% (**Table 1**).

### 3.2 Kinematic Error at Peak Load

The robot reported kinematic error increased when using Sensor Fusion as opposed to Traditional control (**Tables 3 & 4**). However, the motion capture measurements, which don’t

assume robot, fixtures, and bones have no compliance, report decreased error when using Sensor Fusion.

The initial LCL phantom length at the time of digitization was approximately 51 mm and over the course of the varus torque tests the phantom was stretched approximately 4 mm. While the robot reports 0 microns of LCL length error while using Traditional control, the motion capture data reports that there is in fact 101 microns of error after removing the phantom and moving the joint back to the same position (**Table 4**). When utilizing Sensor Fusion for control, motion capture reports a 99% decrease in LCL length error from 101 µm down to 1 µm.

**Table 4.**
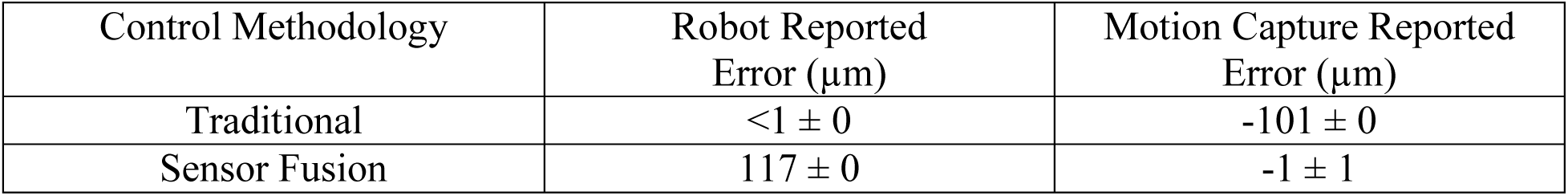
Mean ± SD of the Resultant LCL Phantom Length Error between LCL Phantom Intact and Deficient Conditions at Peak Load.

| Control Methodology | Robot Reported Error ($\mu\text{m}$) | Motion Capture Reported Error ($\mu\text{m}$) |
| --- | --- | --- |
| Traditional | $<1 \pm 0$ | $-101 \pm 0$ |
| Sensor Fusion | $117 \pm 0$ | $-1 \pm 1$ |

### 3.3 Tension and Kinematic Error in Dynamic Loading

Similar to the kinematic error at peak load (**Table 4**), Sensor Fusion control again minimized errors as measured by motion capture (**Table 5**). Here when using Sensor Fusion control, motion capture reports an 89% decrease in RMS error from 61 µm to 7 µm.

**Table 5:** Mean ± SD of the Resultant LCL Phantom Length RMSE between LCL Phantom Intact and Deficient Conditions during Varus Loading & Unloading.

| Control Methodology | Robot Reported RMSE ($\mu\text{m}$) | Motion Capture Reported RMSE ($\mu\text{m}$) |
| --- | --- | --- |
| Traditional | $1 \pm 0$ | $61 \pm 1$ |
| Sensor Fusion | $68 \pm 1$ | $7 \pm 0$ |

In Traditional testing, errors in superposition computed tensions and LCL Phantom length scale with the load applied to the test system (**Figure 7**). These errors are relatively insensitive to applied load using the Sensor Fusion control method. In both Traditional and Sensor Fusion tests, the error is a function of whether the system is being loaded or unloaded. In addition, the path dependent loading/unloading curve shape suggests that there are factors, other than compliance, that can influence tension error.

**Figure 7.**
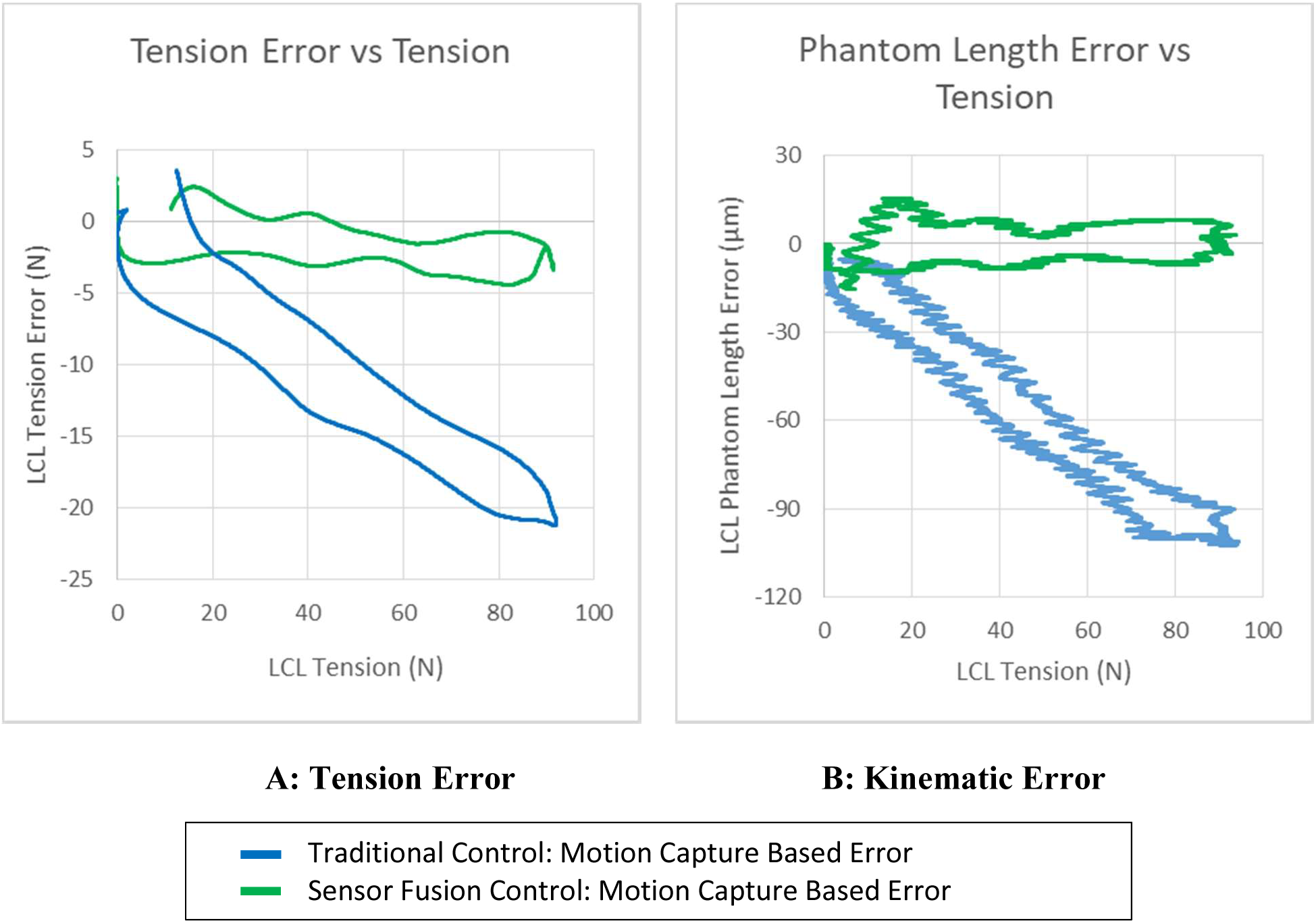
The Position Control LCL tension error (A) and LCL phantom length error (B) relationship with system load. In (A), the single axis load cell data was low pass filtered at 0.2 Hz, similar to the 6-DOF load cell data prior to computing the position control LCL tension error. Filtering was performed in order to remove kinetic noise and make the LCL tension error’s dependence on the loading direction more clear.

### 3.4 Testing Control

According to the joint kinematics measured using motion capture data (**Table 6**), the Sensor Fusion control improved tracking of the prescribed trajectory in the LCL intact and deficient cases. Average RMSE on resultant translation was reduced from 96 µm down to 20 µm. Varus rotation control also improved by about a factor of 3. Robot reported error increased with Sensor Fusion control because the robot end effector was moved differently between intact and deficient surgical conditions to account for the varying compliance the robot, fixtures and bones.

**Table 6:**
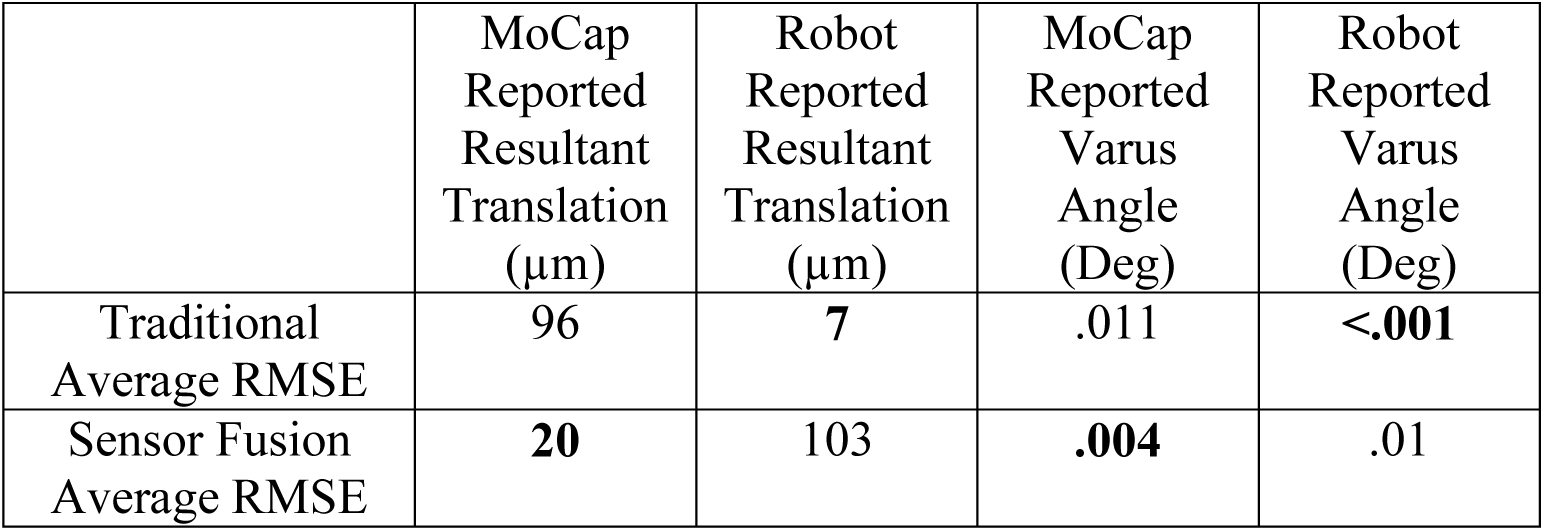
JCS Pose RMSE throughout superposition testing. Metrics used for control are bold.

| | MoCap<br>Reported<br>Resultant<br>Translation<br>($\mu\text{m}$) | Robot<br>Reported<br>Resultant<br>Translation<br>($\mu\text{m}$) | MoCap<br>Reported<br>Varus<br>Angle<br>(Deg) | Robot<br>Reported<br>Varus<br>Angle<br>(Deg) |
| --- | --- | --- | --- | --- |
| Traditional<br>Average RMSE | 96 | 7 | .011 | **<.001** |
| Sensor Fusion<br>Average RMSE | **20** | 103 | **.004** | .01 |

## 4 DISCUSSION

There were 4 key findings from this study. 1) Errors in superposition-computed ligament tensions were reduced from 23% down to 3% using a Sensor Fusion control method compared to using the Traditional control method. 2) Kinematic errors between intact and deficient loading scenarios were reduced by 89%-99% using a Sensor Fusion control method as opposed to a Traditional control method. 3) In addition to test system compliance, there are other loading/unloading path dependent factors that are a source of superposition-computed ligament tension error. 4) Kinematic setpoint tracking errors were reduced 3-4 fold using a Sensor Fusion control method compared to using the Traditional control method.

### 4.1 Tension Error at Peak Load

The first key finding was that the Sensor Fusion control method decreased tension error from 23% down to 3% at peak load. Because the magnitude of system compliance was small (∼101 µm, **Table 4**), this reduction in error in superposition computed tension highlights the importance of accounting for any and all system compliance during superposition testing. These results validate that the Sensor Fusion control methodology works on the surrogate knee and is a promising methodology for use in cadaveric testing. These validation results point to the fact that for several decades there has been a systematic error in superposition testing where system displacement was controlled rather than joint displacement [2]. Researchers have been aware of this error for some time and there have been efforts to mitigate it to some degree by maximizing robot stiffness [2, 25–30], but many researchers use commercially available robots that are not designed for maximal stiffness. This systemic error has likely led to a general underestimation of ligament tension and stiffness [13] and Figure 4 supports these claims. To make matters worse, it should also be considered that when superposition testing has been conducted to understand the contribution of multiple ligaments through sequential cutting, this systemic error may be compounded with each cut [6, 11, 12].

While using a Sensor Fusion control method reduced the errors associated with Traditional superposition testing, it is not a panacea for all test system error. There are most likely several reasons why there is still a 3% position control tension error present when using the Sensor Fusion methodology. One of these is the accuracy of the Optitrak Prime^X^ 13 cameras themselves, which are rated for only ± 200 µm according to Optitrak specifications. Previous work has shown that with filtering and much fine tuning of camera setup parameters, these errors can be reduced considerably, but are nonetheless present. A second source of tension error is the positioning accuracy and resolution of the robot, which inherently has backlash present in its gears. Backlash is explored in more detail in section 4.4. A third source of tension error is that control parameters, such as gains, may cause the system to behave differently when a system’s mechanical properties change, such as when the LCL is intact versus deficient.

When using Sensor Fusion the *force control tension error* is 5% (**Table 1**) while the *position control tension error* is slightly smaller at 3%. This discrepancy is likely due to two reasons. First, robotic force control is generally noisier than position control. This noise shows up in the output kinematic data, which must then be minimized by filtering prior to use as position setpoints for position control trajectories. This filtering can obscure kinematic features within the force control trial. Second, force control gains were used for the intact case (black line in **Figure 6**) but then switched to kinematic gains for the deficient case (blue & red lines), resulting in differing control characteristics. When comparing two position control tests, as in the *position control tension error*, both the position control setpoints and the control gains and other control parameters are more similar, resulting in greater repeatability and lower error.

While the *force control* and *position control tension errors* are similar in this experiment, it is still recommended to do an intact kinematic control test because (1) it shouldn’t add much time to the total testing and (2) the *force control tension error* could be higher if poor decisions for filtering or gains are made.

### 4.2 Kinematic Error at Peak Load

Large tension errors can be caused by very small kinematic errors of 100 µm or less (**Tables 3-5**). In the Traditional control methodology the robot invariably reports no error in kinematics between intact and deficient position controlled tests. If the robot, fixtures, and bones were infinitely rigid then the assumptions of superposition would be satisfied. However, examining the motion capture based data it is clear that the surrogate joint did not go back to the same pose between intact and deficient tests when doing Traditional control. It is only by switching to Sensor Fusion control that motion capture based error was reduced because the algorithm successfully commanded the robot to go to a different pose between intact and deficient test cases to account for changing system compliance.

**Tables 3** & **5** show that the motion capture system measured the Sensor Fusion kinematic error as less than 10 µm. While this is extremely small, actual JCS pose displacements were probably below the noise floor of the motion capture system, so caution should be used when interpreting this error. Regardless, as joint stiffness increases (e.g., as in a joint with a cobalt-chrome implant), similar magnitude displacement errors will result in a larger force errors [13]. Because of this, it is likely that the effect the Sensor Fusion control method would have on tension error will increase in an in-vitro specimen setting because the surrogate knee used in this study was more compliant in compression and varus rotation than a cadaveric knee. To further minimize displacement errors, future test systems could also use 6-DOF position sensors with lower noise floors or higher stiffness robots.

### 4.3 Tension and Kinematic Error in Dynamic Loading

The dynamic error given in **Table 5** demonstrates the fourth key finding, that Sensor Fusion control decreases LCL Phantom length error by 89% in dynamic loading scenarios. **Table 4** showed that in static loading scenarios this decrease was even larger, at 99%. The relatively lower reduction in error during dynamic loading may be due to the difficulties in robotic control during dynamic scenarios.

The Traditional control methodology has a kinematic error, and therefore a tension error, that scales with system load (**Figure 7**) and this is likely due to the system compliance. However, when using Sensor Fusion control, system load and tension error become decoupled. The LCL Phantom length error can be thought of as the amount of kinematic error caused by deflection of the test system, i.e. robot, fixtures, and bone compliance. Therefore, examining the relationship between LCL tension and LCL Phantom length error in Traditional control, it is possible to estimate stiffness of a test system. When examining the same system using Sensor Fusion, the LCL Phantom error doesn’t scale with LCL tension. To be clear, Sensor Fusion doesn’t change the physical stiffness of a test system, but it does gives low stiffness test systems a much higher effective stiffness because it actuates the system to compensate for its own compliance induced error. The Sensor Fusion approach to biological joint testing decreases the negative effects of robot, fixtures, and bone compliance in superposition testing.

Additionally, joint stiffness rather than test system stiffness will still be very important when evaluating test systems using any control method. As stated above, as the joint stiffness increases, smaller kinematic errors will create larger kinetic error. Future work will look at extending this testing methodology to stiffer biological joints. Sensor Fusion control parameters will need to be optimized when dealing with biological tissue rather than a surrogate knee joint with relatively low stiffness. Some of these control parameters include Primary Sensor versus motion capture sensor weighting, filtering, and motion capture setup.

Figure 7 also shows that error is dependent on the direction of loading. While it is tempting to explain this path dependence as the hysteresis of the LCL phantom, these plots are not depicting force vs displacement of the phantom but rather force vs error. For example, the tension error (4) was computed by subtracting the force data from two sensors throughout both loading and unloading (Figure 4). In theory, any differences in force due to loading or unloading would cancel since both force sensors would measure the same force change. The third key finding was that in addition to test system compliance, there are other loading/unloading path dependent factors that are a source of superposition-computed ligament tension error. It is unknown precisely what these factors are, but the error changes sharply when the direction of the robot changes. The green line in Figure 7B demonstrates the phenomena most intuitively, with kinematic error being relatively constant except for when the loading direction switches direction and therefore the kinematic error switches sign. It is hypothesized that the kinematic error could therefore be caused by hysteresis of the force or position sensors themselves, friction within the test system, and/or backlash within the robot.

### 4.4 Testing Control

The fourth key finding was that the Sensor Fusion control method reduced kinematic tracking errors by 3-4 fold. This suggests that the system was able to achieve the target positions, in spite of system compliance. In addition to the applicability of the Sensor Fusion algorithm in superposition testing, this control technique could be applied to other kinematic control processes. Any process where there is significant system load and a need for 6-DOF position or force control with high precision and low bias could benefit from Sensor Fusion approaches. For example, in experiments with viscoelastic effects Sensor Fusion position control can help maintain a prescribed joint displacement rate at high loads even as the robot, fixtures and bones deflect. In experiments with large bone deflection and high loads, Sensor Fusion force control can guarantee that forces are transformed and controlled in the dynamically changing reference frames.

Future work will examine other applications of Sensor Fusion control such as the ability to control specimens of more than 6-DOF. For example, in spine testing each functional spinal unit (FSU) has 6-DOF and they are stacked in sequence. In current testing when there is more than one FSU, additional FSUs are assumed to be 1 rigid body in order to continue to use Traditional control techniques. This assumption limits loading protocols to only pure moments so that all FSU’s experience the same load. By controlling JCS pose based on motion capture sensors mounted close to the articulating surfaces, it is possible to control a single FSU in the middle of a multi-level spinal construct. Similarly force, rather than position, can also be transformed dynamically to that FSU in real time and shear forces can begin to be controlled within a multilevel construct.

## 5 CONCLUSIONS

To **develop** a mechanical testing methodology that corrects for system compliance, sensors familiar to those in the uniaxial testing world (e.g. extensometers), were combined with the concept of visual servoing from robotics literature. A kinematic chain equation (2) was developed which utilizes the high precision, high bias, reliable, and responsive robot encoder data present in Traditional control methods and fuses this with the low precision, low bias, less reliable, and less responsive motion capture markers [19]. By fusing these two sensor outputs together, a more accurate testing methodology that is made that compensates for fixture compliance and directly controls joint motion.

To ***demonstrate*** the effectiveness of the Sensor Fusion methodology, 6-DOF position sensors were mounted to a surrogate knee joint as close as possible to its proximal and distal articulating surfaces. Superposition testing was performed using Traditional testing methods and the Sensor Fusion method. LCL phantom ligament tension was directly measured and compared to superposition-computed tension in the surrogate knee joint. The Traditional control method underestimated ligament tension by 23% while the Sensor Fusion control method brought that error down to 3%.

By conducting superposition testing with additional 6-DOF position sensor(s) in the test system control loop, the commonly overlooked historical accuracy challenges of superposition testing can be greatly reduced. Ultimately, better estimates of ligament contribution to passive joint stability should enable researchers and device manufacturers to better design and refine medical devices and surgical techniques in the future. These new designs and refinements can lead to improvement in clinical outcomes for patients.

## NOMENCLATURE

*_W1_*: World 1: The global static coordinate system that the position of all sensors other than the robot are expressed in. Typically this is the motion capture coordinate system.
*_W2_*: World 2: The static coordinate system of the robot where it is mounted and its positions are reported relative to.
*_P1_*: Primary Sensor 1: The sensor attached to the base rigid body. Typically this is World 1.
*_P10_*: Initial Primary Sensor 2: The relative position of the sensor attached to rigid body 1 when first digitized.
*P2*: Primary Sensor 2: The dynamic coordinate system that is the position of the end of the robot as reported by the robot’s encoders.
*P20*: Initial Primary Sensor 2: The relative position of the sensor attached to rigid body 2 when first digitized.
RB1: Rigid Body 1: The coordinate system of the tool or object mounted to the end of the robot. In the context of cadaveric testing this is usually a bone.
RB2: Rigid Body 2: The coordinate system of the base or object mounted to the world that the tool is interacting with
*JCS Pose*: Displacement of the Rigid Body 2 (RB2 or bone 2) with respect to Rigid Body 1 (RB1 or bone 1) from its neutral position due to an applied load is considered its JCS (Joint Coordinate System) pose. It consists of 3 translations and 3 rotations.
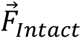: Force measured by the 6-Axis load cell in **Figure 3** with the LCL phantom intact.
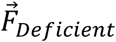: Force measured by the 6-Axis load cell in **Figure 3** with the LCL phantom removed.
𝐿𝐶𝐿_SP_: Lateral collateral ligament phantom tension as measured by the single axis load cell in line with the phantom.
𝐿𝐶𝐿_SA_: Lateral collateral ligament phantom tension as computed by superposition.
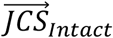: JCS Pose with the ligament intact.
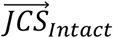: JCS Pose with the ligament removed.
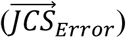: The difference between the intact and deficient JCS Poses.

